# Extracellular matrix mineralization in the mouse osteoblast-like cell line MC3T3-E1 is regulated by actin cytoskeleton reorganization and non-protein molecules secreted from the cells themselves

**DOI:** 10.1101/551531

**Authors:** Hiraku Suzuki, Kazuaki Tatei, Noriyasu Ohshima, Seiichi Sato, Takashi Izumi

**Affiliations:** Center for Medical Education, Graduate School of Medicine, Gunma University, Maebashi, Gunma, Japan; Department of Biochemistry, Graduate School of Medicine, Gunma University, Maebashi, Gunma, Japan; Division of Signaling in Cancer and Immunology, Institute for Genetic Medicine, Hokkaido University, Sapporo, Japan

**Author notes:** Corresponding author: (HS).

## Abstract

Bone tissue constantly undergoes turnover via bone formation by osteoblasts and bone resorption by osteoclasts. This process enables bone to maintain its overall shape while altering its local structure. However, the detailed mechanism of how osteoblast cell-signaling systems induce various structural changes in bone tissue have not yet been completely elucidated. In this study, we focused on the actin cytoskeleton as a regulatory system for bone formation and constructed an *in vitro* experimental system using the mouse osteoblast-like cell line MC3T3-E1. We found that, in MC3T3-E1 cells, the actin cytoskeleton had an important role in matrix mineralization via activation of specific developmental pathways and it was regulated by non-protein molecules secreted from MC3T3-E1 cells themselves. In MC3T3-E1 cells, we observed changes of actin cytoskeleton reorganization and accumulation of PIP_2_ related to actin filament convergences during cell differentiation, in the undifferentiated, early, middle and late stage. Actin cytoskeleton disruption with Cyto D, polymerization inhibitor of actin filament, in early and middle stage cells induced significant increase of osteocalcin mRNA expression normally expressed only in late stage, decrease of Alkaline phosphatase mRNA expression after 24h and abnormal matrix mineralization in MC3T3-E1 cells. Inhibition of Giα with PTX known to regulate actin cytoskeleton in middle stage induced changes in the actin cytoskeleton and PIP_2_ accumulation and suppression of matrix mineralization after 5 days. Furthermore, addition of non-protein molecules from culture medium of cells at various differentiation stage induced difference of PIP_2_ accumulation after 5 min, actin cytoskeleton in 20 min, and matrix mineralization after 5 days. These results not only provide new knowledge about the actin cytoskeleton function in bone-forming cells, but also suggest that cell signaling via non-protein molecules such as lipids plays important roles in bone formation.

## Introduction

Bone tissue consists of cells originating from bone marrow [1, 2], as well as bone matrix [1, 2, 3], which is produced by these cells. Bone matrix is mainly composed of secreted proteins and hydroxyapatite (HA: Ca_10_(PO_4_)_6_(OH)_2_), a type of calcium phosphate. Bone tissue cells can be roughly divided into osteoblasts (bone-forming cells), osteoclasts (which are responsible cell type for bone resorption) and osteocytes (the most abundant type of cell in mature bone-tissue). Balancing bone resorption and production enables bones to both maintain their overall shape and make local structural changes. This process is called bone remodeling [1, 3, 4], and is responsible for bone growth in vertebrates. Disruption of this balance leads to metabolic bone diseases, including osteoporosis [5, 6]. The resorption/production balance plays a role in local changes in bone shape such as sclerosis at sites of inflammation [7] and resorption of mandibular bone in edentulous patients [8, 9]. Growth and injury are known to be the factors causing structural changes of bone. In orthodontics, age of patients [10, 11] and orthodontic force [12, 13] are important elements. However, it is unclear whether bone growth and bone remodeling induced by extracellular stimulation are regulated by each independent mechanisms or by the same system [13]. Because of the structural differences between bone tissue and other organs, and the fact that there are no well-defined functional units consisting of various cell types in bone tissue, unlike in other organs, the complex details of the mechanisms regulating bone growth are largely unknown. However, osteoblasts and osteoclasts interact directly with cell surface proteins that are part of the RANK-RANKL signaling pathway, and this process regulates bone mass by inducing osteoclast differentiation [14, 15]. Osteoclast signaling through this pathway requires RANKL expression on osteoblast cells [15]. Thus, the key to the mechanism of bone regulation must be osteoblasts.

Osteoblasts are not a single cell type, but rather a collection of various bone-forming cell types that have different functional and morphological characteristics, and are sometimes referred to as osteoblast lineage cells. They originate in the bone marrow as mesenchymal stromal cells (MSCs) and gradually differentiate into bone matrix. Osteoblasts work as a cluster, rather than functioning individually [1, 2]. It is unknown how the cell mixture, which secretes a variety of proteins, expresses multiple receptors, and has varying morphological characteristics, integrates its functions [2, 5, 16]. As cell-cell interactions between born-forming cells play an important role in bone mass regulation, in this study we focused on the actin cytoskeleton, which is associated with the regulation of osteoblast form and function [17] and has recently received attention for its role in mechanosensing [17, 18, 19]. We constructed an experimental system based on the mouse osteoblastic cell line MC3T3-E1, and observed actin cytoskeleton functions and changes during cell differentiation in a closed system.

Our results show that the actin cytoskeleton shifts autonomously during MC3T3-E1 cell differentiation, and that stimulation of this process can induce the activation of a specific developmental pathway and remarkable changes in matrix mineralization, which represent the terminal stage of the osteoblast lifecycle [28, 29]. Bone formation results from HA crystal formation and growth in extracellular scaffold proteins, including collagen I, which are regulated by a variety of functional proteins that are secreted from the cells at various stages of differentiation [2, 5]. These functional proteins including Alkaline phosphatase (ALP), which is essential for the initiation of skeletal formation [20] and is regulated by various developmental pathways that interact with each other [1, 4, 21, 22]. Induction of extracellular matrix mineralization in response to one type of stimulation indicated that a specific mechanism activates various developmental signaling pathways and regulates bone formation in response to this type of stimulation. Thus, our results indicate that the actin cytoskeleton is related to basic processes involved in bone formation and cell differentiation. Furthermore, we show that non-protein molecules secreted by MC3T3-E1 cells induce actin cytoskeleton reorganization and changes in matrix mineralization. Although the molecules were not identified, there is some evidence that they are lipids.

This seems likely in light of a recent report indicating the effect of lipids on alveolar bone resorption in periodontal disease [23]. Lipids mediators can be produced quickly from the plasma membrane in an autocrine/paracrine manner [24, 25], and are related to inflammation [26, 27]. Thus, our results provide new information about the actin cytoskeleton and cell signal transduction.

## Materials and methods

### Cell line and culture

MC3T3-E1 osteoblastic cell line was obtained from RIKEN Cell Bank (Tsukuba, Japan), and maintained in Minimum Essential Medium Eagle (αMEM, Sigma-Aldrich, Inc.) containing 10% fetal bovine serum (FBS, Invitorogen), 50 U/ml penicillin, and 50 mg/ml streptomycin. Addition of L (+)-ascorbic acid (AA, Wako) and β-glycerophosphate (βGP, Calbiochem) to the culture medium enables the cells proliferated to the confluence to differentiation and bone formation (Fig. 1F).

**Fig 1.**
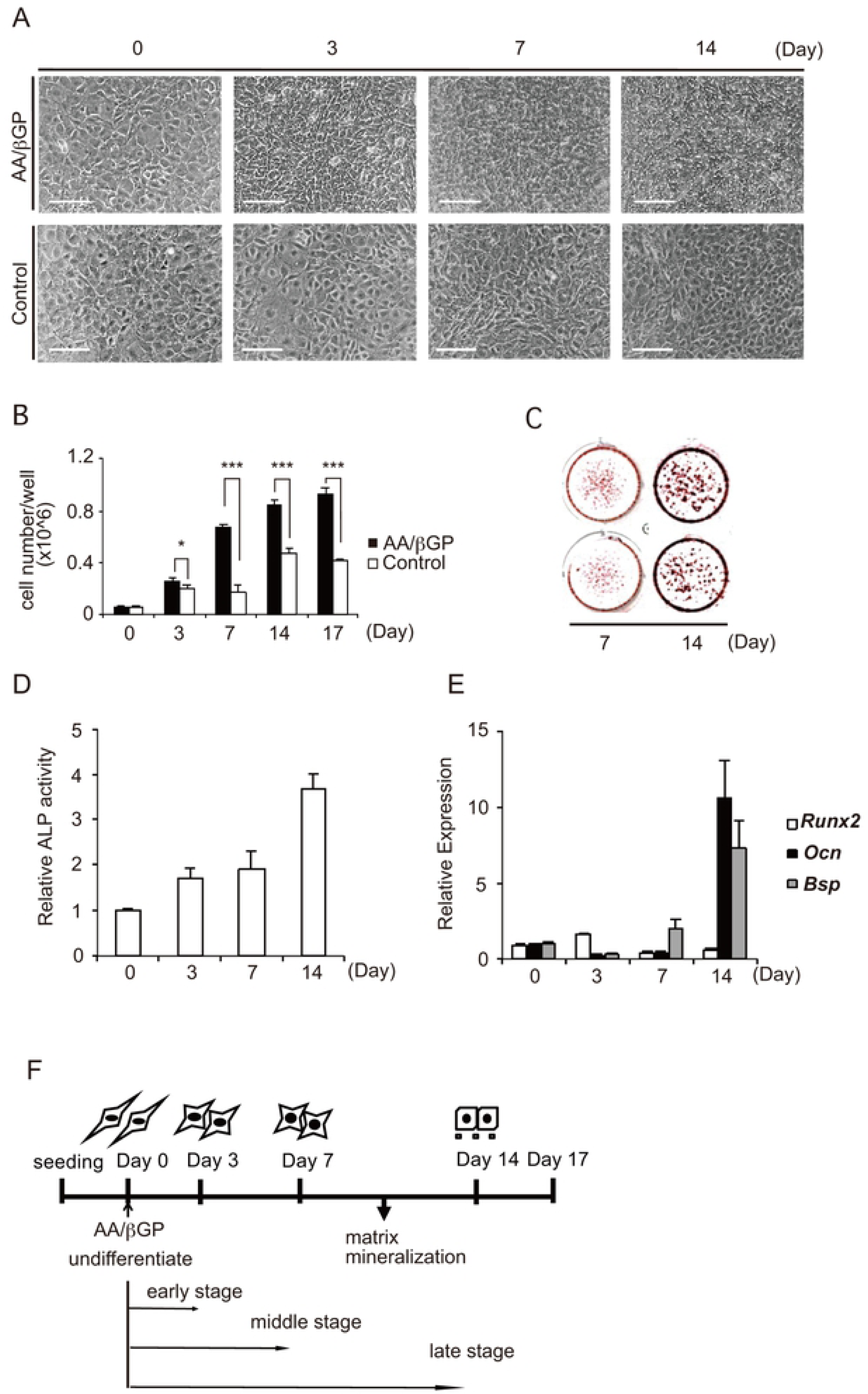
MC3T3-E1 cells differentiation and morphological change is induced by AA/βGP. MC3T3-E1 cells were cultured with or without AA/βGP and used for experiments at the indicated days. (**A**) The cells subjected to morphological change were photographed at 20×magnification at the indicated days. Scale Bar, 100 μm. (**B**) The numbers of cells in a well subjected to culture medium condition were determined at the indicated days. The average data from three independent experiments was expressed as mean ± SD. (**C**) The cells subjected to matrix mineralization were stained by Alizarin Red S at the indicated days. (**D**) Total cell lysates cultured in the presence of AA/βGP were isolated at the indicated days. ALP activities were assayed with the protocol as described in the Material and Methods Section. ALP activity from three independent experiments was expressed as mean ± SD. (**E**) Total RNAs cultured in the presence of AA/βGP were isolated at the indicated days. Quantitative RT-PCR analysis of *Runx2*, *Bsp* and *Ocn* mRNAs as differentiation marker in MC3T3-E1 cells. Relative mRNAs expressions of them were normalized to the expression of *Hrpt1*. (**F**) Assay time course and differentiation stages of our MC3T3-E1 cells are shown. Data are presented as mean ± SD and are representative of at least three independent experiments. Significant difference is expressed as \**P* < 0.05, \*\**P* < 0.01, or \*\*\**P* < 0.001.

### Quantitative RT-PCR

Total RNAs were isolated from culture cells using ISOGENⅡ (Nippon G ene) and were treated with DNAse (TaKaRa) and Recombinant RNAse i nhibitor (TaKaRa). cDNA was prepared from total RNAs using ReverTra Ace (TOYOBO). qRT-PCR was performed using iTaq Universal SYBR Green Supermix (BIORAD) and analyzed on StepOnePlus real-time PCR sy stem (Applied Bionsystems). Data were normalized to the expression of *Hrpt1* for each sample. The primer pairs used for PCR are as follows: *A lp*, forward 5’-ACCTTCTCTCCTCCATCCCT-3’, reverse 5’-GTGTGTGTGTG TGTCCTGTC-3’; *Runx2*, forward 5’-GCCCAGGCGTATTTCAGATG-3’, rever se 5’-GGTAAAGGTGGCTGGGTAGT-3’; *Osx*, forward 5’-TATGCTCCGACCT CCTCAAC-3’, reverse 5’-AATAGGATTGGGAAGCAGAAAG-3’; *Bsp*, forward 5’-GAGGCAGAGAACTCCACACT-3’, reverse 5’-TCTTCCTCGTCGCTTTCCT-3’; *Ocn*, forward 5’-GCAGAACAGACAAGTCCCAC-3’, reverse 5’-ACCTTATT GCCCTCCTGCTT-3’; *Atf4*, forward 5’-TGGCGAGTGTAAGGAGCTAGAAA-3’, reverse 5’-TCTTCCCCCTTGCCTTACG-3’; *Hrpt1*, forward 5’-CAGTCCCA GCGTCGTGATTAG-3’, reverse 5’-AAACACTTTTTCCAAATCCTCGG-3’.

### Alkaline phosphatase activity

ALP activity was measured using p-nitrophenol phosphate method with LabAssay^TM^ ALP (Wako). Whole cell lysate in a well was homogenized in the assay buffer (3.35mM p-nitrophenol phosphate, 1mM MgCl_2_, 50mM Carbonate Buffer/pH 9.8). After 15 min incubation at 37°C, reaction was terminated by the addition of sodium hydroxide solution. The absorbance of p-nitrophenol liberated in the reactive solution was read at 405 nm.

### Alizarin Red S staining

Bone Matrix mineralization was visualized by alizarin red S staining. Briefly, cells were fixed 10% formaldehyde/PBS for 20 min at 4°C after washing with Ca^2+^-free phosphate-buffered saline (PBS) for three times. After five times washing with distilled water, the cells were stained in 1% alizarin red S solution (Wako) for 5 min. The remaining dye was rinsed several times with distilled water. Stained cultures were photographed.

### Immunofluorescence microscopy

MC3T3-E1 cells were plated on 8well microscope slide (SCS-N08; MATSUNAMI) and incubated at 37°C for a specified number of days. The cells were fixed with 3.7% formaldehyde at room temperature (RT) for 10 min. Following washing with PBS, cells were permeabilized by incubation in 0.1% (wt/vol) Triton x-100 (Wako) and blocked by incubation in 1% (wt/vol) bovine serum albumin (A6003; Sigma) in PBS (buffer 1). For PIP_2_ staining, cells were incubated at 4°C overnight in primary antibody anti-PIP2 antibody (ab2335; abcam) diluted in buffer 1 and following washing with PBS, the glass slide was incubated at RT for 1 h in secondary antibody conjugated with Alexa Fluor 488 (A11017; Life Technologies) and SYTOX Orange Nucleic Acid Stain (S-11368; Inviterogen) diluted in buffer 1. For actin cytoskeleton staining, cells were incubated at RT for 1 h in phalloidin conjugated with Alexa Fluor 546 (A22283; Life Technologies) diluted in buffer 1. Subsequently, the glass slide was mounted in 50% (wt/vol) glycerol (Wako) and observed under 40x microscope. Relative fluorescence intensity (PIP_2_/Cell) was calculated from SYTOX-stained nuclei using Image J (RRID: SCR_003070). The images are representative of independent experiments performed in triplicate.

### Non-protein molecules extraction from culture medium

Non-protein molecules secreted from MC3T3-E1 cells were extracted from culture medium of Day 0, Day 3 and Day 7 cell. Except for Day 0, it contained AA and βGP. We incubated the cells with new culture medium under 37°C, 5% CO_2_ for 6h and collected the medium and extracted non-protein molecules using two protocols, Extract 1 and Extract 2.

Protocol 1 (Extract 1): Heat denaturation and centrifugal separation. We heated the culture mediums at 100°C for 5 min. After centrifugal separation at 15,000 rpm for 15 min at 4°C, the supernatants were collected. It mainly includes polar molecules and peptides.

Protocol 2 (Extract 2): Bligh and Dyer method. We added chloroform, methanol and water into the collected medium. The volume of extraction solvents was adjusted the chloroform:methanol:aqueous solution ratio at 1:2:0.8 to single-phase mixture. After vortex, we added chloroform, methanol and water into the solution and adjust the ratio of solution to 1:1:0.9. Centrifugation at 2,500 rpm, 10min at room temperature enabled the solutions separation of aqueous layer and chloroform layer. The organic phase including non-polar molecules like lipids was collected and dried under a stream of N_2_ gas.

After diluting Extract 1s and Extract 2s with culture medium in an appropriate amount, used for the experiments.

### Statistical analysis

Statistical analyses were performed with Microsoft Excel. Values were expressed as means ± SD of triplicate independent experiments. The value was divided by the average value of the controls and the control value set to 1. The difference between the control and experimental groups were compared using relative value. Statistical significance was assumed for \**P* < 0.05, \*\**P* < 0.01, or \*\*\**P* < 0.001.

## Results

### MC3T3-E1 cells differentiation and morphologic changes are induced in ascorbic acid and β-glycerophosphate containing culture medium

Differentiation of osteoblasts and MC3T3-E1 cells can be induced by the combination of ascorbic acid (AA) and β-glycerophosphate (βGP) *in vitro* [28, 29]. In osteoblasts differentiation, not only expression of functional proteins related to bone formation, including ALP, Runx2, bone sialoprotein (Bsp), and osteocalcin (Ocn) [2, 3], but also the morphological features of the cells change during the process of differentiation [2, 16]. We set as Day 0 as 3 days after cell seeding, and cultured the cells for the specified period in the presence or absence of AA/βGP (Fig. 1F). MC3T3-E1 cells cultured in the presence of AA/βGP were closely packed and decreased in size from Day 0 to Day 14, whereas cells cultured without AA/βGP showed little change in size until Day 7 (Fig. 1A). Further, there was a significant difference in cell numbers between the two groups. The number of cells cultured in the presence of AA/βGP increased greatly during the culture period; and while the number of cells cultured without AA/βGP was slightly increased after 7 days of incubation, at Day 14 and Day 17 there were significantly fewer cells compared with those cultured with AA/βGP (Fig.1B). In osteoblasts differentiation, HA crystal formation and matrix mineralization are indicators of ossification and can be visualized by Alizarin red S staining [29]. Calcified nodules resembling woven bone [28] were produced between Day 7 and Day 14 only in the cells cultured with AA/βGP (Fig. 1C). ALP activity of the cells rose continuously from Day 0 to Day 14 (Fig. 1D). Furthermore, we investigated developmental signals that occur during MC3T3-E1 differentiation. We then performed time course analysis of mRNA expression by qPCR after treatment with AA/βGP to determine the mRNA levels of various differentiation markers (Fig. 1E). While the expression of *Runx2*, an early stage differentiation marker, increased at Day 3 but decreased at Day 7, expression of *Ocn* and *Bsp*, which are middle to late stage differentiation markers, increased from Day 7 to Day 14 and reached a peak of approximately 10 times the original expression level. These results indicated that the culture with AA/βGP induced MC3T3-E1 differentiation. Thus, we succeeded in constructing an experimental system that integrates the function and form of bone-forming cell clusters, and which can be used to observe how the cells respond to different stimuli at various stages of differentiation. We concluded that MC3T3-E1 cells were undifferentiated at Day 0 and in the early, middle, and late stages of differentiation on Day 3, Day 7, and Day 14, respectively (Fig. 1F).

### Actin cytoskeleton reorganization occurs during MC3T3-E1 cell differentiation

The dynamics of the actin cytoskeleton, which is formed by F-actin microfilaments, are related to cell differentiation [16, 17]. In MC3T3-E1 cells, the actin cytoskeleton plays an important role in cell migration [18] and responds to external force by forming stress fibers [18, 19]. Using a method described in previous reports, we stained F-actin with Alexa Fluor 546-labelled phalloidin [30, 31] and observed the cytoskeleton during MC3T3-E1 cell differentiation (Fig. 2A). On Day 0, abundant actin filaments were observed in the cells, and they decreased in number by Day 3, Day 7, and Day 14. On Day 7, the F-actin had assembled and appeared as thick, bright fibers. However, by Day 14, the fibers were beginning to unravel and exhibited many cracks. Next, we observed changes in Phosphatidylinositol 4, 5-bisphosphate (PIP_2_) localization associated with cell differentiation by Alexa Fluor fluorescent antibody staining [32]. PIP_2_ is a phospholipid that is located in the cell membrane and is a known substrate for a number of important signaling proteins and is related to actin fiber assembly [33–37]. Cytosolic PIP_2_ levels were significantly higher in MC3T3-E1 cells on Day 0 and Day 7 compared with Day 3 and Day 14 (Fig. 2B). Quantitative assessment based on the ratio of PIP_2_ fluorescence intensity to cell number (as determined by nuclear staining with SYTOX) showed that the PIP_2_/cell number value was significantly higher on Day 0 and 7 compared with Day 3 and Day 14 (p < 0.01) (Fig. 2C). However, there was no difference in actin fiber formation or PIP_2_ localization in undifferentiated MC3T3-E1 (data not shown). Thus, we concluded that actin cytoskeleton dynamics are related to differentiation and bone-forming capacity in MC3T3-E1 cells.

**Fig 2.**
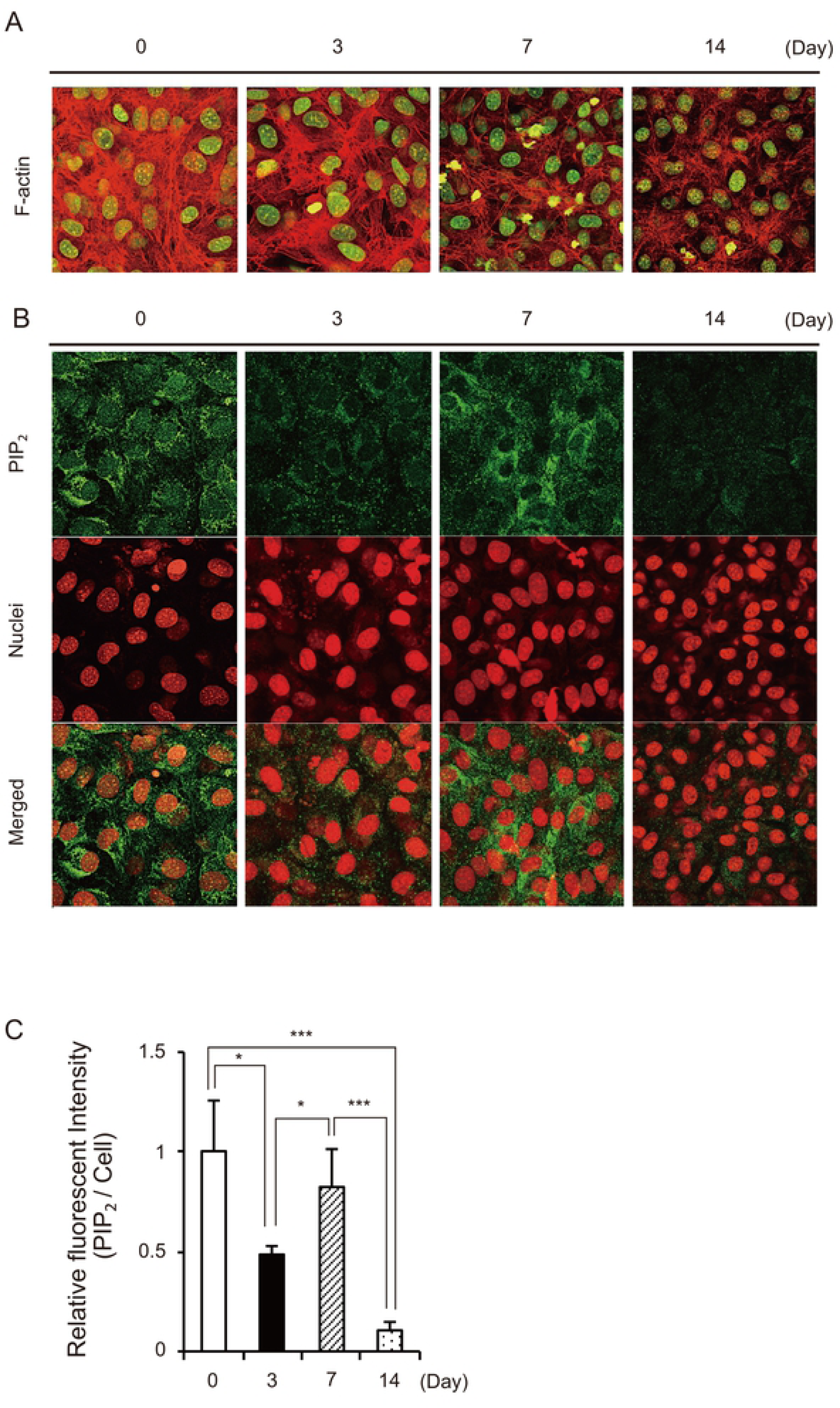
Differentiation of MC3T3-E1 cells induces actin cytoskeleton reorganization and the change of PIP_2_ localization. MC3T3-E1 cells were cultured with AA/βGP and used for experiments at the indicated days. (**A**) The cells subjected to actin cytoskeleton reorganization were stained with Alexa Fluor 546-labeled phalloidin for F-actin of actin filaments (red) and SYTOX-green for nucleuses (green). (**B, C**) The cells subjected to PIP_2_ localization were stained with anti-PIP_2_ antibody and Alexa fluor 488 for PIP_2_ localization (green) and SYTOX-orange for nucleuses (red) (B). Fluorescent intensity ratios of PIP_2_ to MC3T3-E1 cell in (B) were shown. Data are presented as mean ± SD and are representative of at least three independent experiments (C). Significant difference is expressed as \**P* < 0.05 or \*\*\**P* < 0.001.

### Disruption of actin filaments with cytochalasin D inhibits normal matrix mineralization by MC3T3-E1 cells

Previous investigations have shown that cytoskeletal reorganization and cell morphology play an important role in MC3T3-E1 differentiation. To elucidate the functional role of actin cytoskeleton in matrix mineralization, we treated MC3T3-E1 cells with the cell permeable fungal toxin cytochalasin D (Cyto D, Sigma). This compound binds to the barbed end of actin filaments and inhibits actin polymerization [30, 38]. Incubation of MC3T3-E1 cells with Ctyo D at 37°C for 1 h on Day 3 disrupted actin filaments (Fig. 3A), and the same results were obtained on Day 7 and Day 14 (data not shown). To investigate whether disruption of the actin cytoskeleton affected MC3T3-E1 cell matrix mineralization, we added Cyto D to the MC3T3-E1 cell culture medium at Day 3, Day 7, and Day 14. After 3 days, we changed the medium and continued to culture the cells until Day 17, when Alizarin red S staining was performed. Treatment with 2 μM Cyto D on Day 3 and Day 7 significantly inhibited normal matrix mineralization and calcified nodule formation; however, treatment on Day 14 has no influence on matrix mineralization (Fig. 3B). In enlarged images of MC3T3-E1 cells cultured under normal conditions, we observed some extracellular HA crystals that stained with Alizarin red S, whereas cells that were incubated with Cyto D did not exhibit these crystals; only a type of structure around the cells and the stained cells themselves were seen (Fig. 3C). Taken together, these results indicate that there is relationship with between actin cytoskeleton and matrix mineralization in MC3T3-E1 cells.

**Fig 3.**
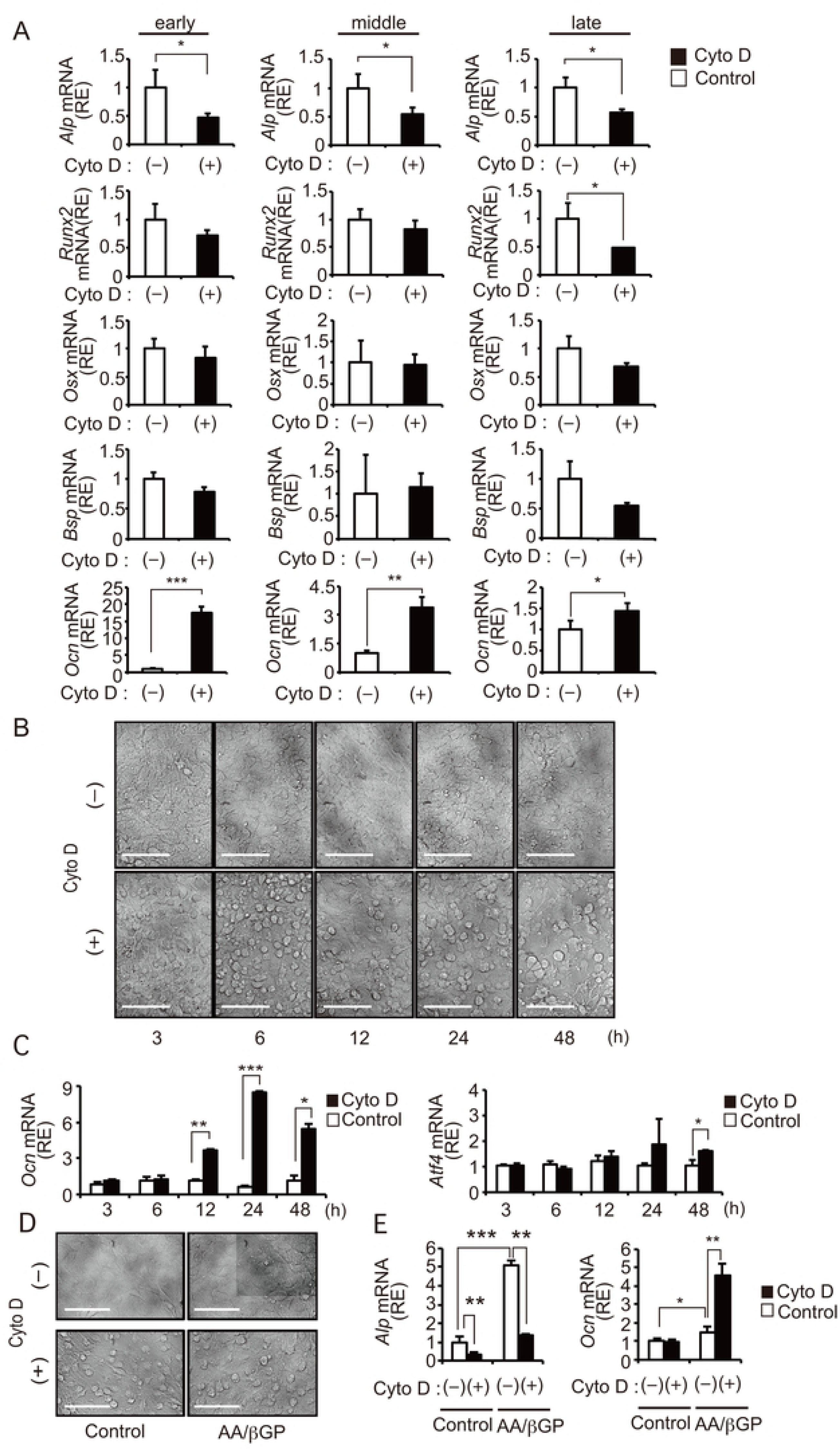
Actin cytoskeleton disruption with Cyto D inhibited normal matrix mineralization of MC3T3-E1 cells. MC3T3-E1 cells were cultured with AA/βGP and stimulated with Cyto D at the indicated differentiation stages (Day 3: early, Day 7: middle, Day 14: late stage). (**A**) The cells subjected to actin cytoskeleton disruption at early stage were stimulated with Cyto D (2 μM) for 1 h and stained with Alexa Fluor 546-labeled phalloidin for F-actin of actin filaments (red) and SYTOX-green for nucleuses (green). (**B**, **C**) The cells subjected to matrix mineralization were stimulated with Cyto D (2 μM) for 72 h at the indicated stages and stained with Alizarin red S at Day 17 (B). Light microscopic examinations (40×) of the cells stimulated with Cyto D (2 μM) at early stage and stained with Alizarin red S at Day 17 (C). Left is images with a focus on cells, right is images with a focus on Alizarin red S stain (Bar: 100 μm).

### Cyto D induced actin cytoskeleton disruption activates specific developmental pathways associated with MC3T3-E1 differentiation

Matrix mineralization is regulated by many developmental pathways in bone-forming cells [2, 4, 5]. The failure in HA crystal formation observed in MC3T3-E1 cells incubated with Cyto D in the early and middle stages of differentiation indicated that Cyto D inhibited early developmental signals, including regulation of ALP, which plays an important role in extracellular HA crystal formation and growth [20]. To determine which developmental pathway is affected by Cyto D, we measured changes in mRNA expression by RT-PCR. Twenty-four hours after incubation with Cyto D, there was a significant decrease in the mRNA expression level of the early stage differentiation marker *Alp*, and increased mRNA levels of the late stage differentiation marker *Ocn* were detected at all stages of MC3T3-E1 differentiation; however, the most remarkable change in mRNA expression was observed in early stage cells (Fig. 4A). In contrast, mRNA expression levels of the early stage differentiation markers *Runx2* and *Osx* and the late stage differentiation marker *Bsp* did not change. ALP and osteocalcin expression are affected by Wnt signaling pathway activation, and their expression is inversely related to osteoblast differentiation [2, 21, 22]. Cyto D induced morphological changes in early stage MC3T3-E1 cells that could be observed at 3 h and became marked after 6 h (Fig. 4B). A significant increase in *Ocn* mRNA levels occurred 12 h to 48 h after stimulation (Fig. 4C left), and peaked at 24 h. Furthermore, mRNA expression of the multiple transcription factor *Atf4* [39] increases significantly 48 h after treatment with Cyto D (Fig. 4C right). Even though marked changes in cell morphology and significant decrease in *Alp* mRNA expression were observed in both cells cultured with/without AA/βGP for 3 days (Fig. 4D, 4E), a significant increase in *Ocn* mRNA expression was observed only in cell cultured in the presence of AA/βGP (Fig. 4E). These results show that Cyto D induced actin cytoskeleton disruption and cell shape change induced specific developmental pathways related to *Alp* and *Ocn* mRNA expression. It suggests that actin cytoskeleton reorganization associate with cell differentiation of MC3T3-E1 cells.

**Fig 4.**
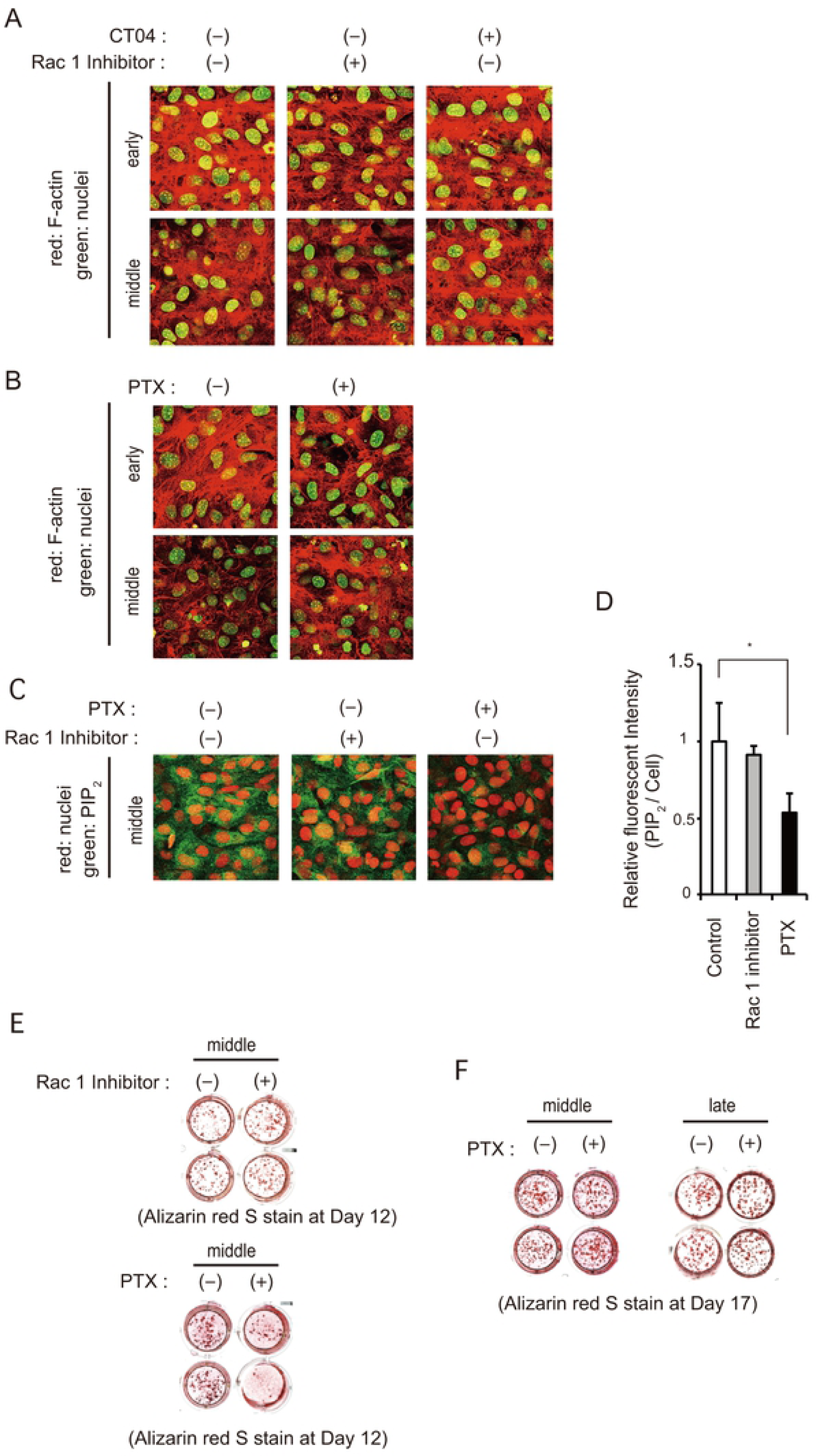
Cyto D induces morphologic change and activation of a specific developmental pathway in MC3T3-E1 cells. MC3T3-E1 cells were cultured with or without AA/βGP and stimulated with Cyto D at the indicated differentiation stages (Day 3: early, Day 7: middle, Day 14: late stage). (**A**) Quantitative RT-PCR analysis of *Alp*, *Runx2*, *Osx*, *Bsp* and *Ocn*mRNAs after 24 h of stimulation with Cyto D (2 μM) in MC3T3-E1 cells at the indicated differentiation stages. Relative mRNAs expression of them were normalized to the expression of *Hrpt1*. Data are presented as mean ± SD and are representative of at least three independent experiments. (**B, C**) The cells were cultured with AA/βGP and stimulated with Cyto D (2 μM) at early stage for 3, 6, 12, 24, 48 h. Light microscopic examinations (40×) are shown (Bar: 100 μm) (B). At 3, 6, 9, 12, 24, 48 h after stimulation, total RNAs were subjected to qRT-PCR analysis for *Ocn* (left) and *Atf4* (right). Data were normalized to the expression of *Hrpt1* (C). **(D, E)** The cells were cultured with or without AA/βGP for 3 day and stimulated with Cyto D (2 μM) for 24 h. Light microscopic examinations (40×) are shown (Bar: 100 μm) (D). At 24 h after stimulation, total RNAs were subjected to qRT-PCR analysis for *Alp* (left) and *Ocn* (right). Data were normalized to the expression of *Hrpt1* (E). Significant difference is expressed as \**P* < 0.05, \*\**P* < 0.01, or \*\*\**P* < 0.001.

### Actin cytoskeleton reorganization associated with MC3T3-E1 differentiation is regulated via the Rac1 signaling pathway

Our results showing that actin cytoskeleton function is associated with matrix mineralization are consistent with a previous report [40]. The actin cytoskeleton is involved in cell adhesion, cell migration, and stress fiber formation [30, 40, 43] in bone-forming cells. Its function is regulated by the Rho family of G proteins, including Cdc42, Rac1, and RhoA [40, 41, 42, 43]. To identify which actin cytoskeleton regulation pathway is related to MC3T3-E1 matrix mineralization during cell differentiation, we inhibit Rac1 and Rho function in MC3T3-E1 cells in the early and middle stages of differentiation using selective Rac1 inhibitor (Wako) and Rho inhibitor (CT04: Cytoskeleton, Inc). After incubation with the Rac1 inhibitor at 37°C for 4 h, there was a clear decrease in the number of actin filaments visible in MC3T3-E1 cells on Day 3 and a slight decrease on Day 7 (Fig. 5A). In contrast, no cytoskeletal differences were observed in MC3T3-E1 cells incubated with the Rho inhibitor compared with the control. Our results are consistent with a previous report that Rac1 inhibition influence osteoblast differentiation [41, 42]. However, there were differences in the effects of Rac1 GEF on actin cytoskeleton between early stage and middle stage MC3T3-E1 cells. This may imply that, in addition to Rac1, actin cytoskeleton reorganization is also regulated by other factors that participate in actin filament assembly.

**Fig. 5.**
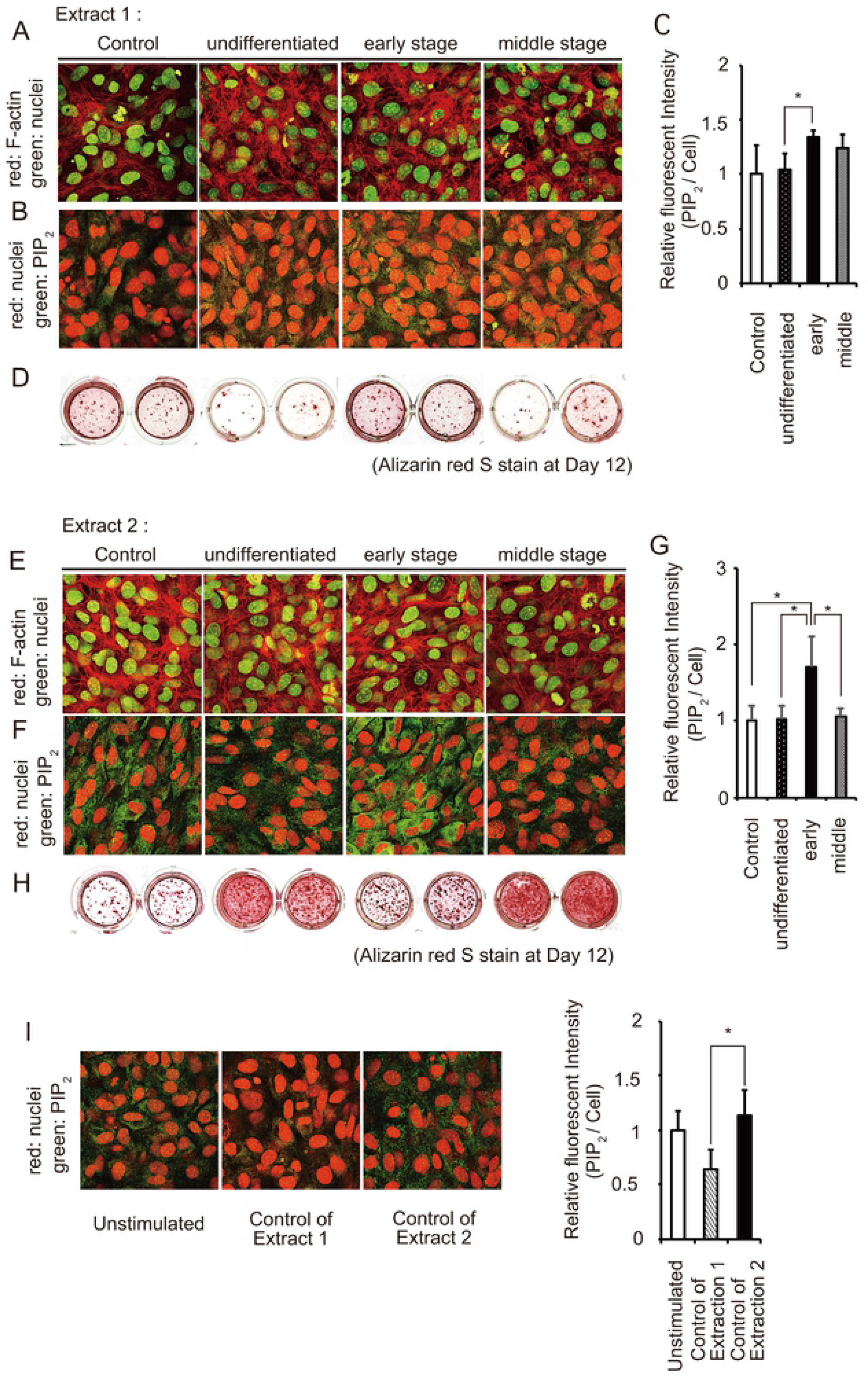
Actin cytoskeleton reorganization and matrix mineralization of MC3T3-E1 cells are regulated via GPCR pathway. MC3T3-E1 cells were cultured with AA/βGP and stimulated with the reagents at the indicated differentiation stages (Day 3: early, Day 7: middle, Day 14: late stage). (**A**) The cells subjected to actin cytoskeleton reorganization at the indicated stages were stimulated with Rho Inhibitor (C3 exoenzyme:2.0 μg/ml) or Rac1 Inhibitor (50 μM) for 4 h and stained with Alexa Fluor 546-labeled phalloidin for F-actin of actin filaments (red) and SYTOX-green for nucleuses (green). (**B**) The cells subjected to actin cytoskeleton reorganization at the indicated stages were stimulated with PTX (1.0 μg/ml) for 4 h and stained with Alexa Fluor 546-labeled phalloidin for F-actin of actin filaments (red) and SYTOX-green for nucleuses (green). (**C**, **D**) The cells subjected to PIP_2_ localization at middle stage were stimulated with PTX (1.0 μg/ml) for 4 h and stained with anti-PIP_2_ antibody and Alexa fluor 488 for PIP_2_ localization (green) and SYTOX-orange for nucleuses (red) (C). Fluorescent intensity ratios of PIP_2_ to MC3T3-E1 cell in (C) were shown. Data are presented as mean ± SD and are representative of at least three independent experiments (D). (**E**) The cells subjected to matrix mineralization were stimulated with Rac1 Inhibitor (50 μM) (up) or PTX (1.0 μg/ml) (down) for 72 h at middle stage and stained with Alizarin red S at Day 17. (**F**) The cells subjected to matrix mineralization were stimulated with PTX (1.0 μg/ml) for 3 days at middle stage (left) or late stage (right) and stained with Alizarin red S at Day 21. Significant difference is expressed as \**P* < 0.05.

### Actin filament assembly via GPCR regulates MC3T3-E1 cell matrix mineralization in the middle stage of differentiation

Actin cytoskeleton organization in MC3T3-E1 cell is regulated by Rac1 [40–42], which we also confirmed here. PIP_2_ production is regulated by phospholipase C (PLC) and G protein-coupled receptors (GPCRs) [35, 45]. GPCRs play an important role in various signaling pathways, and are reported to be involved in cytoskeletal rearrangement [45]. The Rac1 pathway is also reported to be regulated by GPCRs, as well as Giα [31, 45], which is a heterotrimeric G protein. To examine whether GPCR regulates actin cytoskeleton reorganization in MC3T3-E1 cells, we stimulated MC3T3-E1 cell in the middle stage of differentiation with 1.0 μg/ml pertussis toxin (PTX, Calbiochem) [31]. PTX catalyzes ADP-ribosylation of the α subunit of the Giα protein and prevents its interaction with GPCRs. As a result, incubation with PTX for 4 h at 37°C reduced the number of actin filaments in early stage MC3T3-E1 cells and inhibited filament assembly in middle stage cells (Fig. 5B). Furthermore, stimulation with PTX inhibited PIP_2_ localization (which is associated with cell differentiation) in the cytosol of middle stage cells, whereas treatment with the Rac1 inhibitor did not have a significant influence on PIP_2_ localization (Fig. 5C, D). The Rac1 inhibitor inhibits interactions between Rac1 protein and the Rac-specific guanine nucleotide exchange factors (GEFs) Trio and Tiam1 [41]. We next examined the effects of inhibiting G protein signaling on matrix mineralization, which is required for terminal differentiation of MC3T3-E1 cells. We added the Rac1 inhibitor or PTX to the culture medium in MC3T3-E1 cells in the middle stage of differentiation and incubated them under normal conditions for five days (until Day 12). Matrix mineralization was only suppressed in MC3T3-E1 cells stimulated with PTX (Fig. 5E). However, treatment with PTX during the middle stage of differentiation had no apparent effects by Day 17, and PTX treatment of late stage MC3T3-E1 also had no effect on matrix mineralization by Day 17 (Fig. 5F). These results suggest that actin fiber formation stimulated by the GPCR pathway during the middle stage of MC3T3-E1 cell differentiation regulates matrix mineralization.

### Non-protein molecules secreted by MC3T3-E1 cells regulate actin cytoskeleton reorganization and matrix mineralization of MC3T3-E1 cells themselves

Using our MC3T3-E1 experimental system, we showed that actin cytoskeleton reorganization is associated with matrix mineralization regulation, and that the structure of the cytoskeleton is modified during cell differentiation. PTX affected actin cytoskeleton reorganization, PIP_2_ accumulation, and matrix mineralization in MC3T3-E1 cells in a closed system, indicating that actin cytoskeleton reorganization is regulated by GPCRs and their ligands in an autocrine manner. Although bone-forming cells secrete many auto-regulatory proteins [1, 2, 3], we hypothesized that non-protein molecules may also regulate actin cytoskeleton organization and matrix mineralization. To test this, we collected culture medium from undifferentiated, early and middle stage MC3T3-E1 cells that had been incubated for 6h under normal conditions. We extracted non-protein molecules from the culture media by first performing a 100 °C, 5 min heat denaturation step, followed by centrifugation at 4 °C and 15,000 rpm for 15 min, or via the Bligh-Dyer extraction method, which is widely used in lipid research [46, 47]; the former method isolates including mainly polar molecules and peptides that are dissolved in the culture medium (Extract 1), and the latter method only isolates mainly non-polar molecules (Extract 2). We processed fresh culture medium in the same two ways to serve as controls. We stimulated middle stage MC3T3-E1 cells with Extract 1s and Extract 2s and observed PIP_2_ localization 5 min later, actin cytoskeleton organization 20 min later, and changes in matrix mineralization 5 days later (at Day 12). The cells treated with Ex1 from undifferentiated cells and cells in the middle stage of differentiation exhibited a markedly lower level of matrix mineralization compared with control cells and cells treated with Extract 1 from early stage cells (Fig. 6D). However, only a small difference was observed in PIP_2_ localization and actin cytoskeleton reorganization (Fig. 6A, 6B, 6C). Treatment with Extract 2 from undifferentiated and middle stage cell resulted in a significant decrease in PIP_2_ localization signal (Fig. 6F, 6G) and actin cytoskeleton degradation (Fig. 6E) compared with cells treated with Extract 2 from early stage cells and control. Furthermore, normal matrix mineralization was inhibited like the cells treated with Cyto D in these cells on Day 12 (Fig. 6H, Fig. 3B). Promotion of normal matrix mineralization was observed in the cells treated with Extract 2 from early stage cells (Fig. 6H), which showed significant increase in PIP_2_ localization signal compared to control (Fig. 6F, 6G). Control of Extract 1, including only extraction from αMEM and FBS induced significant decrease of PIP_2_ signal intensity compared to Control of Extract 2 (Fig. 6I). Non-protein molecules extracted with two methods from different cell stages both induced the difference of matrix mineralization on Day 12, however, matrix mineralization changes with actin cytoskeleton reorganization were induced by non-protein molecules in Extract 2. Thus, these results showed that non-protein molecules, especially non-polar molecules secreted from MC3T3-E1 cells regulated matrix mineralization via actin cytoskeleton reorganization.

**Fig 6.**
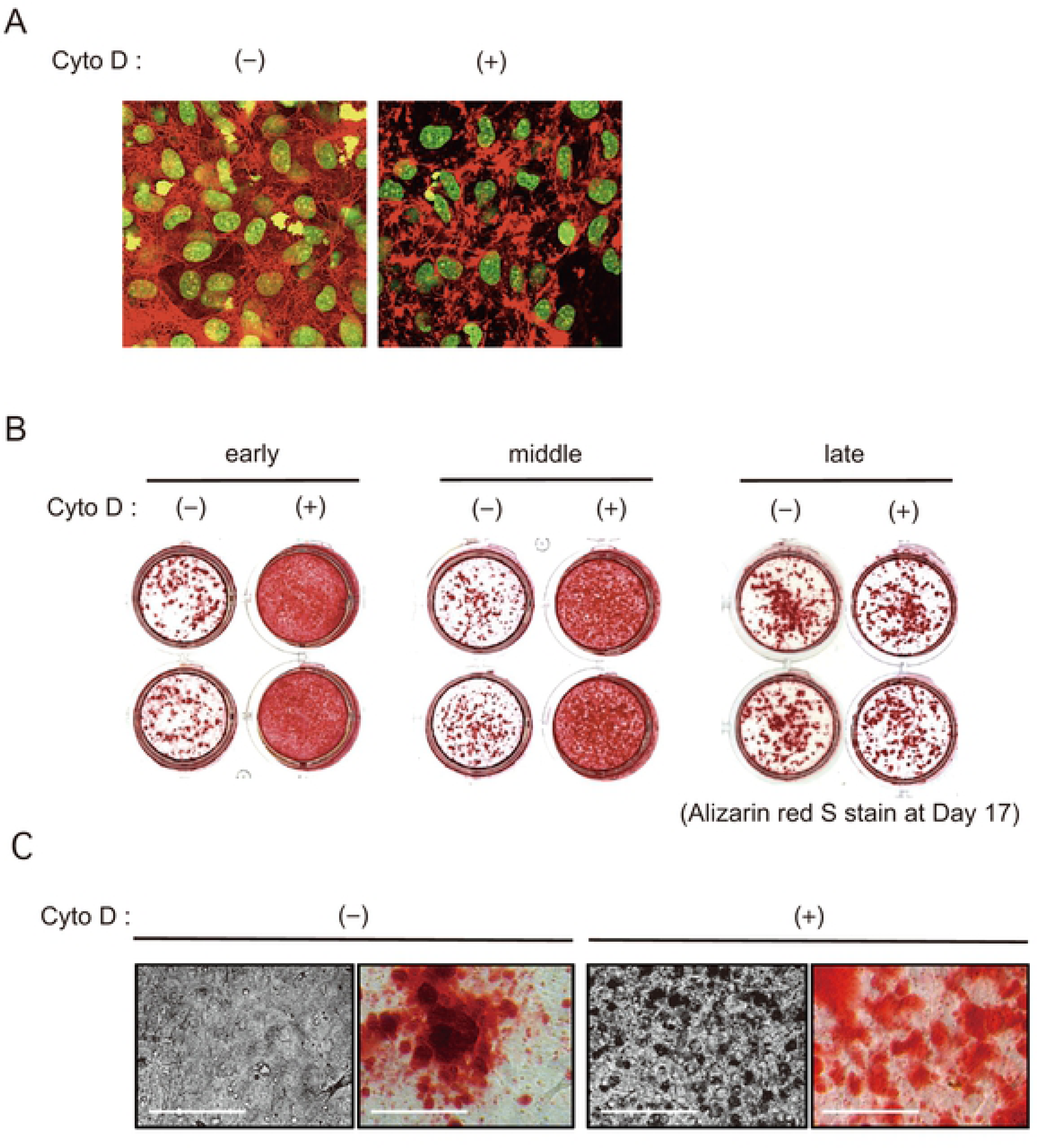
Non-protein molecules secreted from MC3T3-E1 cells induces actin cytoskeleton reorganization and matrix mineralization change. MC3T3-E1 cells were cultured with AA/βGP and used for experiments at the indicated stages. Extract 1s included polar molecules and Extract 2s included non-polar molecules secreted from MC3T3-E1 cells at the indicated stages. (**A**) The cells subjected to actin cytoskeleton reorganization at the middle stages were stimulated with Extract 1 for 20 min and stained with Alexa Fluor 546-labeled phalloidin for F-actin of actin filaments (red) and SYTOX-green for nucleuses (green). (**B, C**) The cells subjected to PIP_2_ localization at middle stage were stimulated with Extract 1 for 5 min and stained with anti-PIP_2_ antibody and Alexa fluor 488 for PIP_2_ localization (green) and SYTOX-orange for nucleuses (red) (B). Fluorescent intensity ratios of PIP_2_ to MC3T3-E1 cell in (B) were shown. The results were representative of at least three independent experiments. Data represent mean ± SD (C). (**D**) The cells subjected to matrix mineralization were stimulated with Extract 1 for 72 h at middle stage and stained with Alizarin red S at Day 17. (**E**) The cells subjected to actin cytoskeleton reorganization at the middle stages were stimulated with Extract 2 for 20 min and stained with Alexa Fluor 546-labeled phalloidin for F-actin of actin filaments (red) and SYTOX-green for nucleuses (green). (**F, G**) The cells subjected to PIP_2_ localization at middle stage were stimulated with Extract 2 for 5 min and stained with anti-PIP_2_ antibody and Alexa fluor 488 for PIP_2_ localization (green) and SYTOX-orange for nucleuses (red) (F). Fluorescent intensity ratios of PIP_2_ to MC3T3-E1 cell in (G) were shown. The results were representative of at least three independent experiments. Data represent mean ± SD (G). (**H**) The cells subjected to matrix mineralization were stimulated with Extract 2 for 72 h at middle stage and stained with Alizarin red S at Day 17. (**I**) The cells subjected to PIP_2_ localization at middle stage were stimulated with control of Extract 1 or Extract2 for 5 min and stained with anti-PIP_2_ antibody and Alexa fluor 488 for PIP_2_ localization (green) and SYTOX-orange for nucleuses (red) (left). Fluorescent intensity ratios of PIP_2_ to MC3T3-E1 cell were shown (right). The results were representative of at least three independent experiments. Data represent mean ± SD. Significant difference is expressed as \**P* < 0.05.

## Discussion

Bone remodeling occurs via osteoblast-mediated bone formation and osteoclast-mediated bone resorption. Resorption of jaw bone tissue in edentulous patients [8, 9] indicates that mechanical stimulus plays an important role in maintaining proper bone shape and structure. In this case, bone resorption proceeds at a relatively moderate pace, whereas extreme bone resorption and sclerosis are observed at injured or inflamed sites [7]. Thus, we predicted there were multiple pathways to detect and manage chronic and acute extracellular stimulation in bone tissue, in addition to well-known bone remodeling system. The actin cytoskeleton detects extracellular stress [17] and has recently been reported to help regulate bone tissue maintenance [19, 30]. We hypothesized that the regulatory mechanism we were searching for involved the actin cytoskeleton; however, to observe the actin cytoskeleton in bone-forming cells, we need to first solve a few problems. First, careful consideration was required to determine how to evaluate the effect of a single activating signal on bone formation, as bone-forming cells are regulated by many developmental pathways. Second, in bone-forming cells, activation of developmental pathways was inhibited by each other during cell differentiation [21, 22]. Thus, we had to construct an experimental system in which we could minimize the effects to woven bone formation derived from interaction of bone-forming cells in various differentiation stages.

First, we used the osteoblast-like cell line MC3T3-E1 to construct an experimental system with standardized bone-forming cells. MC3T3-E1 cells can be induced to differentiate using a combination of AA and βGP [28, 29], so MC3T3-E1 cells in a single culture plate well can function as an independent cell cluster. In our study, MC3T3-E1 cells incubated in the presence of AA and βGP exhibited changes in differentiation marker mRNA expression levels (Fig. 1E), ALP activation (Fig. 1D), calcified nodule formation (indicating matrix mineralization) (Fig. 1C), cell morphology, and the number of cells (Fig. 1A, B), depending on the culture day. mRNA expression of *Runx2*, a master regulator of bone formation and early stage differentiation marker, was up-regulated during Day 0-3, and down-regulated during Day 7-14. Bone sialoprotein (BSP) and osteocalcin (OC) functioned in matrix mineralization are known as late stage differentiation markers. The expression of *Bsp* and *Ocn* mRNA increased notably during Day 7-14, consistent with calcified nodule formation (Fig. 1C). These results indicate that different lengths of incubation with AA and βGP promoted changes in cell differentiation stages. The MC3T3-E1 cell clusters that formed by Days 0, 3, 7, and 14 can be considered to resemble preosteoblasts, immature osteoblasts, mature osteoblasts, and osteocytes, respectively (Fig. 1F). Because MC3T3-E1 cell cluster differentiation was induced in a closed system, we concluded that certain molecules secreted by MC3T3-E1 cells themselves promoted differentiation marker expression and stimulated the growth of these cells in an autocrine manner. Thus, we constructed an experimental system containing bone-forming cells in which we were able to regulate the differentiation stage.

Next, we observed the changes in actin cytoskeleton organization during MC3T3-E1 cell differentiation in our experimental system. The actin cytoskeleton plays a role in mechanosensing [17, 19, 30], cell adhesion [17, 40, 43], and migration [17, 43]. The number of actin filaments in MC3T3-E1 cells gradually decreased over time and degraded at Day 14 (Fig. 2A). PIP_2_ localization, which is associated with actin filament binding [35], diminished temporarily at Day 3, increased at Day 7, and disappeared at Day 14 (Fig. 2B, C). As a similar transient change was observed in *Runx2* mRNA expression (Fig. 1E), we hypothesized that actin cytoskeleton reorganization was associated with cell differentiation. Differentiated osteoblasts turn into osteocytes that are individually buried in bone matrix [1, 2, 3]. Presumably, the degradation of the actin cytoskeleton that was observed during MC3T3-E1 cell differentiation was consistent with this loss of stimulus-responding ability and cell migration. These results show that the actin cytoskeleton reorganization during MC3T3-E1 cell differentiation and osteogenesis (Fig. 2A).

Here, we investigated the effects of actin cytoskeleton reorganization on MC3T3-E1 cell differentiation and matrix mineralization. Extracellular matrix mineralization in MC3T3-E1 cells (Fig. 1C) was observed with remarkable increase of *Ocn* mRNA expression in late stage (Fig. 1E). In late stage, actin fiber degradation was seen in MC3T3-E1 cells. In the early and middle stages of MC3T3-E1 cells, addition of Cyto D, which inhibits F-actin polymerization [31, 38], induced disruption of the actin cytoskeleton (Fig. 3A) and resulted abnormal matrix mineralization (Fig. 3B). In the enlarged image, calcified nodules was not observed (Fig. 3C) and significant decrease of *Alp* mRNA expression and remarkable increase of *Ocn* mRNA expression after 24 h in qRT-PCR analysis (Fig. 4A). ALP is essential for initiating skeletal mineralization in the early stage of MC3T3-E1 cell differentiation [20, 29]. Osteocalcin has a high affinity for calcium and plays an important role in HA crystal growth [48, 49, 50]. Thus, we hypothesize that Cyto D-induced abnormal matrix mineralization was due to inhibition of mineralization mediated by suppression of ALP expression and early expression of osteocalcin. It is consistent with the previous report that the presence of large amount of osteocalcin in synovial fluid of rheumatoid arthritis (RA) and osteoarthritis (OA) patients caused skeletal deformity [51]. In our study, *Ocn* mRNA expression was strictly suppressed in the early stage of cell differentiation under normal conditions (Fig. 1E). Expression of ALP and some early stage differentiation marker proteins are regulated by the Wnt/β-catenin pathway, and osteocalcin and other late stage differentiation markers are controlled by the BMP-2 pathway. The activation of Wnt/β-catenin pathway downregulates activation of BMP-2 pathway [21, 22]. In bone tissue, it is assumed that switching of these developmental pathways in osteoblasts differentiation control the quantitative change of ALP responsible for HA crystal formation and osteocalcin related to crystal growth and induce normal matrix mineralization. Treatment with Cyto D in late stage in which *Ocn* mRNA expression increased did not caused abnormal matrix mineralization (Fig. 3B) and induced slight change in *Alp*/*Ocn* mRNA expression (Fig. 4A) were consistent with our hypothesis. Involvement of actin cytoskeleton in cell differentiation was supported by the differences of *Alp*/*Ocn* mRNA expression induced by addition of Cyto D in cells cultured with or without AA/βGP (Fig. 4E). Furthermore, treatment with Cyto D induced an increase of *Atf4* mRNA expression (Fig.4C), known as stress-responsive transcription factor [2, 41], suggested further developmental pathway activation interactions. Thus, our results suggested that the actin cytoskeleton functioned as a trigger for activation of a specific developmental pathway and has an important role in matrix mineralization in bone-forming cells.

Subsequently, we investigated how the actin cytoskeleton reorganization regulated during cell differentiation. The actin cytoskeleton can act as a mechanosensor, regulated by Rho [17, 30, 44], and as a migration and adhesion regulatory factor, modulated by the Rac1 pathway [17, 40, 41]. In our study, we showed the actin cytoskeleton change was induced by treatment with a Rac1 inhibitor in the MC3T3-E1 cell in mainly early stage (Fig. 5A). This indicates that actin cytoskeleton reorganization during cell differentiation was associated with Rac1 independently of mechanical stress. Furthermore, localization changes of PIP_2_ during cell differentiation remind us to activation changes of PLC producing inositol 1, 4, 5-triphosphate (IP_3_) and diacylglycerol (DAG) from PIP_2_. It is reported that PLC function in Wnt signaling pathways [34, 48] and both PLC and Rac1 are associated with GPCR signaling [31, 45, 53] which detect various extracellular stimuli. As we expected, the cells treated with PTX, Giα selective inhibitor, in middle stage showed suppression of assembly of actin filaments (Fig. 5B), PIP_2_ localization (Fig. 5C, D) and matrix mineralization on Day 12 (Fig.5E). In addition, the results that Rac1 inhibitor did not affect PIP_2_ localization and matrix mineralization of MC3T3-E1 cells in middle stage suggested that matrix mineralization regulation with actin cytoskeleton is mainly controlled via GPCR. Moreover, PTX effects on matrix mineralization were observed only in limited period, from Day 7 to Day 12 (Fig. 5F). It indicated that the regulation system of matrix mineralization via GPCR was strongly influenced by differentiation stage of MC3T3-E1 cells.

The results that treatment with PTX caused changes in PIP_2_ localization, actin cytoskeleton and matrix mineralization indicates that MC3T3-E1 cells received some stimulation via GPCRs. Thus, we hypothesized that not only functional proteins but GPCR ligands secreted from MC3T3-E1 cells regulated actin cytoskeleton and subsequent activation of developmental pathways of MC3T3-E1 cells themselves in autocrine/paracrine manner. Some GPCR ligands are lipid mediators, including PGE_2_, 2AG, and S1P, which function in bone-forming cells [27, 54, 55]. Lipids are produced and degraded much more quickly than proteins. Further, second messenger cAMP [56], Ca^2+^ [6] and propeptides [57] are also reported to influence bone formation. If these non-protein molecules regulate reorganization of the actin cytoskeleton in bone-forming cells, then they should have a strong effect on matrix mineralization. We therefore removed proteins that directly regulate matrix mineralization, like ALP, and investigated whether non-protein molecules secreted by MC3T3-E1 cells regulate the actin cytoskeleton and matrix mineralization. From the results and previous reports about GPCRs, we focused on the nature of the molecules secreted from MC3T3-E1 cells and its quantitative and qualitative changes. We collected culture medium from MC3T3-E1 cells in various differentiation stages and extracted non-protein molecules using two methods. Extract 1 mainly included water-soluble molecules in aqueous culture medium, meaning polar molecules including cAMP, Ca ion and peptides. Extract 2, which was extracted by the Bligh & Dyer lipid extraction method [46, 47], contained non-polar molecules, including lipids. As a result, in both Extraction 1 and Extraction2 assay, molecules extracted from different cell stages induced the difference of matrix mineralization on Day 12. A more significant influence on PIP_2_ localization, actin cytoskeleton reorganization and matrix mineralization were observed in Extraction 2 assay (Fig. 6E, 6F, 6G, 6H) whereas, overall suppression of PIP_2_ signal and matrix mineralization was observed in cells treated with Extract 1 compared to the Extract 2 treated cells (Fig. 6A, 6B, 6C, 6D). Effects of Extract 2 indicated that non-protein, non-polar molecules secreted from MC3T3-E1 cells during cell differentiation regulated matrix mineralization via actin cytoskeleton reorganization. Significant difference between control of Extract 1 and control of Extract 2 (Fig. 6I) might be due to polar molecules from FBS, which suppressed PIP_2_ signaling and matrix mineralization. Nevertheless, the difference of matrix mineralization induced by Extract 1 including polar molecules secreted from different cell stages were observed (Fig. 6D). This result suggested that non-protein molecules secreted from MC3T3-E1 also regulated matrix mineralization independent of PIP_2_ signals and it is consistent with previous reports of second messengers [6, 56, 57]. Thus, we showed that non-protein molecules secreted from MC3T3-E1 cells during cell differentiation regulated matrix mineralization via actin cytoskeleton reorganization.

In this study, we showed that the actin cytoskeleton of MC3T3-E1 cells functioned as a trigger of a specific developmental pathway and is a rapid and conclusive matrix regulation factor that is controlled by non-protein molecules secreted by the MC3T3-E1 cells themselves. This suggests that not only functional proteins, but also non-protein molecules, including lipids, have an important role in the regulation of bone formation. Recently, lipid contributions to cell differentiation and bone disease have been reported [23, 24, 27, 53, 54]. Thus, our findings helps to elucidate part of the complex interaction of bone-forming cells and show that we need to consider the influence of these non-protein molecules in bone research and treatment. However, lipids are too unstable to detect with previous methods; in addition, there are many lipids and many GPCRs that bind to them. We are now trying to identification of lipids secreted from MC3T3-E1cels during cell differentiation by mass spectrometry.

Our experimental system is an amplifier system that allowed us to visualize cell responses by eliminating cell-cell interactions between various types of bone-forming cells and limiting cell characteristics. Presumably, the results obtained using this model are different from the responses that occur in actual bone tissue, but it is suitable for observation of both rapid responses and matrix mineralization changes that are mediated by a single stimulus. Not only these findings, but also future work using this experimental system may be helpful for orthodontics, surgical treatment, research, and treatment of bone disease.

## Acknowledgments

We thank Emily Crow, PhD, from Edanz Group (www.edanzediting.com/ac) for editing a draft of this manuscript.

